# The eco-evolution and biodiversity in antagonistic and mutualistic interactive populations

**DOI:** 10.1101/2023.02.01.526361

**Authors:** Lingzi Wang, Mikael Pontarp

## Abstract

Understanding environmental and ecological effects on ecosystem structure, function, and dynamics is crucial. However, our comprehension of such effects remains challenging due to the complex interplay of abiotic environment, ecological relationships among species, and coevolution within and across trophic levels. Here, we set out to investigate a wide range of combined ecoevolutionary responses in a plant-insect ecosystem. We study ecological population dynamics, coevolution and functional diversity in response to changes in (1) abiotic-biotic interactions, (2) within-trophic-level competitive interactions, and (3) between-trophic-level antagonistic and mutualistic interactions. We developed an eco-evolutionary and functional trait-based model to simulate environmental and ecologically mediated responses to these interactions. We show that mutualistic interactions render a positive feedback loop of increased population sizes followed by selection for even stronger biotic ecological interactions instead of abiotic interactions. Antagonistic interactions have the opposite effect and adaptation to abiotic environment becomes more important. Furthermore, our study reveals that benign conditions (high ecological opportunities, low within-trophic-level competition, and strong interactions between plants and insects) facilitate functional diversity, whereas harsh conditions impede diversity. These findings provide valuable insights into mechanisms that underlie fundamental eco-evolutionary responses to current environmental and ecological changes, offering potential contributions to ecosystem conservation facing urgent challenges.

## Introduction

The societal and economic benefits associated with ecosystems and several of the services within are indisputable (Alexandridis et al. 2021; Potts et al. 2016). Simultaneously, our actions are reshaping the environment, ecology, and biodiversity, which have evolved over extensive temporal scales (Miller, 2005). For example, agricultural intensification negatively impacts many insect species by reducing within-field habitat quality, among-field diversity, and the amount of semi-natural habitat (Díaz *et al*. 2019; Wagner *et al*. 2021). These anthropogenic endeavours, often driven by short-term objectives, are disrupting ecosystems and leading to biodiversity loss (Díaz et al., 2006; Hilderbrand et al., 2010). These actions pose a potential threat to the long-term sustainability of robust ecological dynamics and health (Cardinale, 2012; Loreau et al., 2022; Naeem, 2002).

Despite the paramount importance of understanding the maintenance of ecological systems, progress in understanding the processes that underlie ecosystem structure (e.g., biodiversity) and function has however been slow (Loeuille et al., 2013). One reason for the slow progress arguably lies in the complex interplay of environmental effects, ecological interactions and evolutionary processes that shape diverse ecosystems. Organisms engage not only with their abiotic surroundings but also with other biotic populations both within and between-trophic-levels. The ecological consequences of these interactions combined with plausible coevolution imply the importance of eco-evolutionary processes in dynamical ecosystems (Cortez & Ellner, 2010; Ellner et al., 2011; Urban & Skelly, 2006). Yet, understanding the eco-evolutionary processes that generate coevolution and functional diversity in ecosystems that include a complex mix of environmental effects, as well as antagonistic and mutualistic interactions, remain elusive (Pennekamp et al., 2018; Urban et al., 2016). Much remains to be learned regarding how these different factors interact within ecosystems.

With the above being said, previous research provides some theoretical bases for understanding the maintenance and diversification of complex ecosystems. Adaptation in functional traits and adaptive radiations associated with some biotic and abiotic interactions are, considered to reflect the reasons for coevolution and diversification processes driven by eco-evolutionary processes in the ecosystems. For example, studies show that the coevolution and diversity of various organisms can be driven by ecological opportunity, such as diversified habitats or climate conditions (Losos, 2010; Meyer & Kassen, 2007; Stroud & Losos, 2016; Yoder et al., 2010). These varied ecological niche spaces allow the emergence of new species, and the gradual accumulation of genetic and functional differences (Bolnick and Fitzpatrick., 2007; Wellborn & Langerhans, 2015). Another important driver of the partitioning of niche spaces during diversification is the competition pressure for limited resources (e.g., food, territory, or mates). The competition pressure can lead to the emergence of new traits or ecological roles, which foster the coexistence of multiple spaces in the same ecosystem (Dieckmann & Doebeli, 1999; Doebeli & Dieckmann, 2000; Mac Arthur & Levins, 1967, Pontarp et al., 2012, Pontarp et al., 2015, Pontarp et al., 2017). Other studies emphasize the antagonistic and mutualistic trophic interactions as the drivers of adaptation and diversification. Theory shows the coevolution and adaptive radiations in an antagonistic community context (Brännström et al., 2011; Braga, 2021; Ito et al., 2009; Loeuille & Loreau, 2005; Sauterey et al., 2017; Wang et al., 2021;), and in specific predation can induce evolution and disruptive selection on prey populations and drive the evolutionary branching of prey (Brown & Vincent, 1992; Ito et al., 2009; Pontarp & Petchey, 2018; Pontarp, 2021; Ripa et al., 2009; Wang et al., 2019). Furthermore, mutualistic interaction can also affect population abundance (Aguilera et al., 2020), trait evolution (Ramos & Schiestl, 2019) and biodiversity (Aguilera et al., 2020; Häussler, J. et al., 2017). For instance, studies show that mycorrhizal fungi coevolve with different host plants, which can cause the emergence of various fungal species, each specialized for specific plant partners (Brundrett., 2002; Smith & Read., 2008).

Although the above theories focus on different specific aspects (i.e., ecological opportunity, competition, antagonism and mutualism) that have their influences on the ecological function, coevolution or diversification, these aspects likely impact the ecosystem in concert (Vellend 2016). For example, the characteristics of an antagonistic herbivore population may have an indirect impact on a mutualist population even though they do not interact directly with each other, but indirectly through a third population (Ohgushi, 2005). Therefore, it is worth adopting a holistic view of ecosystems as a food web of multiple types of interactions (e.g., Brodersen et al., 2018) and investigating a wide range of combined eco-evolutionary responses. With this in mind, we develop an eco-evolutionary and functional trait-based model to explore environmental and ecologically mediated responses to these interactions. Our model focuses on three main types of interactions that populations commonly face: abiotic-biotic interactions, within-trophic-level competition, and between-trophic-level antagonistic and mutualistic interactions (Figure 1). It follows that we focus on three different interacting trophic levels viewed here as herbivores, pollinators, and plants. We explore the ecological responses (the change of population sizes) and evolutionary responses (coevolution in functional traits and diversification in biological populations with distinct functional traits) to the three types of interactions. We use the adaptive dynamics approach (introduced by Geritz et al., 1998) for the analyses.

**Figure 1.**
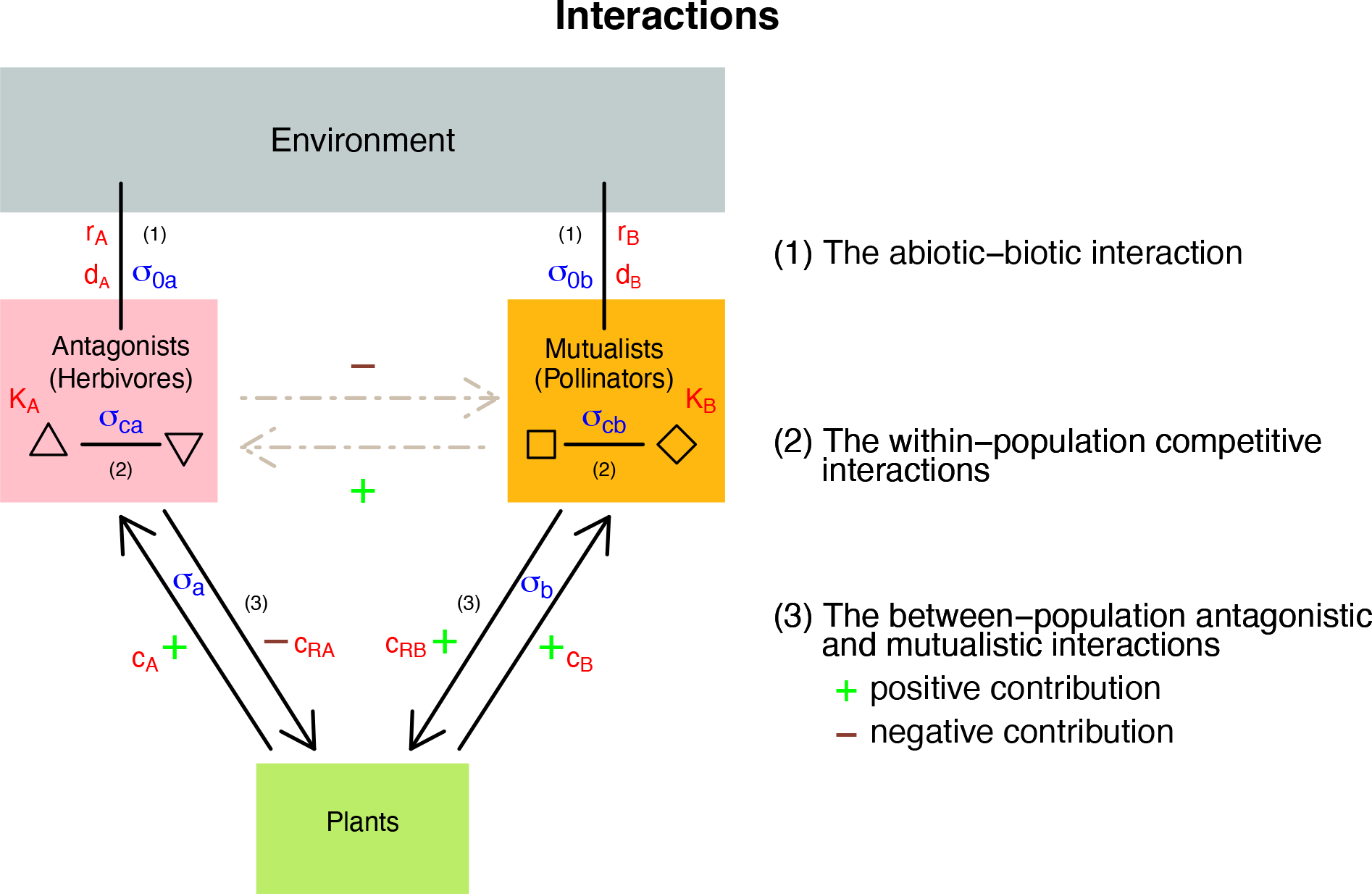
Illustration of the three populations and interactions in the ecological network. Interactions are denoted by numbers (1-3) and represented by black lines/arrows. (1) The strength of abiotic-biotic interactions is represented by the birth rates (*r*_A_, *r*_B_) and the death rates (*d*_A_, *d*_B_) while the antagonists (*A*) and the mutualists (*B*) live in abiotic environment. (2) The strength of within-trophic-level competitive interactions is represented by the carrying capacity of the antagonist and mutualist populations (*K*_A_, *K*_B_) to compete. (3) The strength of between-trophiclevel antagonistic and mutualistic interactions is represented by the corresponding interactive convergent coefficients for trophic-level plant resources (*c*_A_, *c*_RA_, *c*_B_, *c*_RB_). The interactive ranges of the three interactions are measured by the niche widths in the three interactions: σ_0a_, σ_0b_ in the interaction (1); the competition niche width: σ_ca_, σ_cb_ in the interaction (2); and antagonistic and mutualistic interactive niche width: σ_a_, σ_b_ in the interaction (3).

## Methods

### Overview

We focus on three main interactions that most biological populations will face in their ecosystems (1) environmental selection pressures (i.e., environmental interactions), (2) within-trophic-level competitive interactions, and (3) between-trophic-level antagonistic and mutualistic interactions (Figure 1), and explore how these three interactions will affect ecological dynamics, trait coevolution and biodiversity in an interactive ecological network. We view the populations in our system as a set of antagonistic herbivores, mutualistic pollinators, and plants on which the herbivores and pollinators rely (Figure 1). We assume that functional traits in the biological populations drive all the interactions. Such traits may be any traits that are of relevance for environmental adaptation and ecological interactions within or between populations. Body size is, for example, a trait that is of particular relevance as size commonly correlates with many other functional traits including physiology, phenology, feeding morphology, etc. (White et al., 2007). Irrespective of what specific traits are considered we generalized them in a mathematical trait-based model of ecological and eco-evolutionary dynamics (Brännström et al., 2013). More specifically, we generalise traits by assuming that herbivores or pollinator populations possess the traits of relevance both for their adaptation to the environment and their interaction with other populations. We also assume that the optimal trait for the given environment and the trait that maximises resource acquisition from the plants may be different, i.e., assuming a tradeoff between abiotic environmental adaptation and biotic plant resource acquisition. Furthermore, by assuming that traits can evolve, it follows that population sizes and functional traits change dynamically in the system in response to environmental effects, resource availability and ecological interactions. Herbivore and pollinator traits that affect antagonistic and mutualistic interactions in the system evolve dynamically and can also branch and diversify, so the populations with the new traits become new functional populations. The newly emerged distinct functional populations then automatically join the interactions with the existing populations. We run the dynamics until the whole eco-evolution and diversification process reaches the stable equilibrium. To this end, we use the adaptive dynamics approach (Brännström et al., 2013; Dieckmann & Law., 1996; Doebeli & Dieckmann., 2000; Geritz et al., 1998) (details in the section “Eco-evolutionary analysis”).

### Trait-based ecological model

The focused three populations in the model are the plant resource population (*R*), the antagonistic herbivore population (*A*), and the mutualistic pollinator population (*B*). Population interactions are illustrated in Figure 1. In the interaction with abiotic environment (1), both the herbivore and the pollinator populations *A* and *B* are dependent on their adaptation to their environment based on how their functional traits *a* and *b* deviate from the optimal traits for the environment *a*_0_, *b*_0_ and corresponding interactive niche widths σ_0a_, σ_0b._ The strength of the interactions with the environment is represented by their birth rates r_A_, r_B_ and death rates *d*_A_, *d*_B_ (Figure 1). In specific, the degree of adaptation to the environment acts on the growth of the populations by the factor of the normal kernel exp 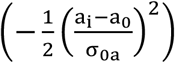 for the herbivores and exp 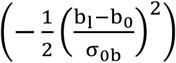 for the pollinators (eq. I) and their respective intrinsic growth rates r_A_ and r_B_. It follows that adaptation to the environment is strongest when the population trait equals an optimal adaptive environmental trait value *a*_0_ or *b*_0_. Also, adaptation tapers off symmetrically as the trait value mismatches the optimal value according to environmental adaptation niche width parameters σ_0a_ and σ_0b_. We thus model adaptation to the general environment using a concept of trait matching between the organismal trait and the optimal trait value for a given environment. We combine such trait matching with the concept of niche width where a wide niche width implies that the organism is a generalist that suffers relatively less from having a mismatched environmental trait while a specialist suffers more from such a mismatch. The growth rate and the death rates of the plant resource *r* and *d* are assumed to be constant and so not affected by the environment (Lotka-Vertera Model, Lotka, 1920; Lotka, 1925; Volterra, 1926).

Both the herbivore and pollinator populations engage in within-trophic-level competitive interactions for their own limited mutual resource, denoted as carrying capacity *K*_A_ and *K*_B_ (Figure 1 interaction (2)). We assume that the competitive interaction among populations depends on the same trait as the ones described above (in interaction (1)) and trait matching between populations dictates competitive strength. Such a trait may again be body size which commonly correlate with other more specific traits of relevance for competition for resource, nesting places, etc (White et al., 2007). Similar to the niche concept presented above we assume that such interactions diminish following the functional form of a normal distribution. More specifically, the strength of the competition acts on population growth by the factor of 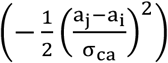and exp 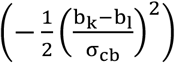 (eq. I), where σ_ca_ and σ_cb_ denote the competition niche width for herbivores and pollinators.

The herbivores and pollinators also have trophic-level interaction with their shared plant resource *R*. The plant population thus have positive effects on both herbivores and pollinators (Figure 1 interaction (3)). The effect from the herbivores on the plants is negative since the herbivores feed on the plants, whereas the effect from the pollinators on the plants is positive due to mutualistic contributions (Figure 1 interaction (3)). Like the interaction strength mentioned above in interactions (1) and (2), the strengths of the interaction between the plant and the herbivore populations depend on the matching between herbivore population trait (*a*_i_), their corresponding optimal interactive trait with the plants (*a*_m_), and also on the resource niche width σ_a_. It follows that the strength of the interaction is highest when the herbivore trait equals the optimal interactive trait with plant trait (*a*_m_), and tapers off when the herbivore and plant trait value mismatches following a normal kernel exp 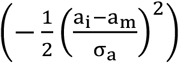. We also introduce the conversion coefficients *c*_A_, *c*_RA_ which denote how much the strength of interaction between the herbivores and the plants are converted into their own growth (eq. I). Similarly, for the positive mutualistic interactions between pollinators and plants, the strength of their interaction follows a normal distribution exp 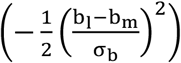, with the conversion coefficient c_B_, c_RB_ for the two populations (eq. I). It is worth noting that the pollinators and the herbivores also have indirect interaction with each other through their interaction with their shared plant resources (dashed arrows in Figure 1).

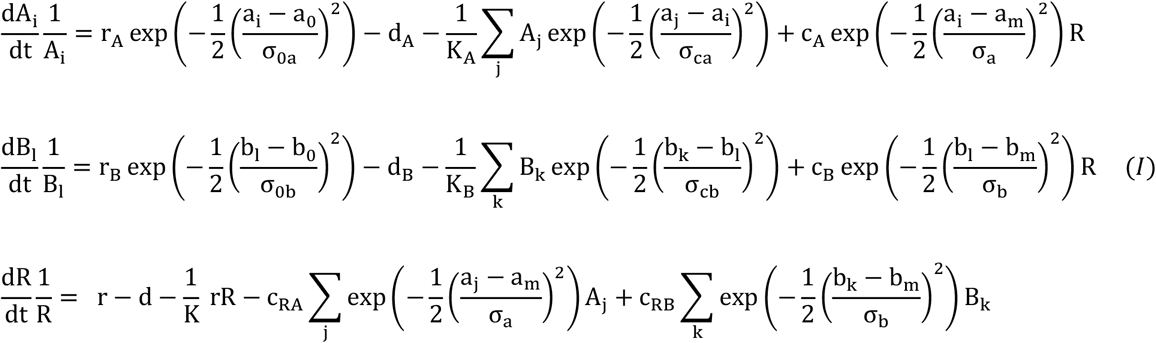

(*i, j* = 1, 2, …, *N* and *l,k* = 1, 2, …, *M*, where *N* and *M* are the total number of populations of herbivores and pollinators)

### Eco-evolutionary model analyses

To investigate the frequency-dependent ecological population dynamics and evolutionary dynamics, we apply adaptive dynamics theory (Brännström et al., 2013; Dieckmann & Law., 1996; Doebeli & Dieckmann., 2000; Geritz et al., 1998). According to the theory, ecological dynamics are assumed to be faster than the evolutionary process, i.e., ecological equilibria are reached before any mutation-driven trait changes are introduced. For the ecological equilibrium, we used computational method (in *R* 4.2.0) to retrieve the ecological population size equilibrium, which happens when all the population sizes do not change (the equilibrium population sizes are also checked as the fitness of all populations being zero using the ‘solve’ function in *R* 4.2.0). At the ecological equilibrium, an evolutionary mutation will occur only when the mutational traits bring a positive fitness value to the focused mutating populations, which follow the idea of “invasion fitness” by Metz et al (1992), i.e. a population with a mutant trait will be able to invade the resident population given that the mutant population have a positive invasion fitness at ecological equilibrium. Evolutionary mutations are assumed to be small, i.e., the mutant trait values replace the nearby resident trait value gradually in the evolutionary process. For such gradual trait evolutionary processes, we use the Canonical equation (Dieckmann & Law., 1996) to calculate the trait changes as a function of time:

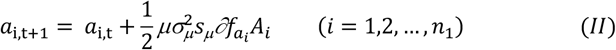

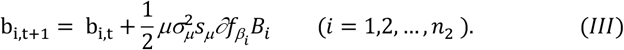

where *a*_i,t_, 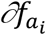 and *A*_*i*_ in Eq. II represent the value of *i*-th antagonist’s trait, selection gradient (the derivative of the fitness value in respect to the trait value) and the population size at the time *t* correspondingly (similarly for the mutualists in Eq. III), and *μ*,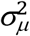, *S*_*μ*_ represent the mutation rate, mutation variance, and mutation steps, which are fixed in our analysis.

Following the above procedure of ecological equilibrium and mutation invasions, novel traits will continuously replace the resident trait until the resident fitness reaches the local maximum or minimum in the fitness landscape (Brännström et al 2013). If a fitness maximum is reached, evolution will stop as no more mutants can invade. However, if the resident trait evolves to the local minimum in the fitness landscape, the fitnesses of mutating traits on both sides of the resident trait are higher than the fitness of the resident trait, so disruptive selection emerges. The population then diverge into two phenotypically distinct populations, and both populations join the interactions with the ecosystem and the evolutionary dynamics continue (Geritz et al., 1998). For example, the herbivore population or the pollinator population can diverge from the original population *A* or *B* into two and later into several more population *A*_i_ (i=1,…,*n*_1_) or *B*_l_ (l=1,…,*n*2) with their corresponding traits *a*_i_ (*i*=1,…,*n*_1_) or b_l_ (*l*=1,…,*n*_2_). The competitive interaction then emerges among all the populations *A*_i_ (*i*=1,…,*n*_1_) and among all the populations *B*_l_ (*l*=1,…,*n*_2_) respectively. The newly generated populations will coevolve with the other existing populations (eq. I). It is also possible that during this coevolutionary process, one or more population sizes drop to zero and thus go extinct. All the populations continue to coevolve with the new diverged populations as well as adapt to new extinctions until all the population trait values reach their local maxima in the fitness landscapes. Under such conditions, when all populations are on peaks in the fitness landscape, the evolutionary dynamics become stable and stop changing, which means that the Evolutionary Stable Strategy has been reached (the *ESS*, Geritz et al., 1998). All results presented below are analyzed at the *ESS*.

### Model scenarios and analyses

Here we use the above eco-evolutionary model to delve into the mechanisms governing ecosystem structure, function and dynamics, focusing on the impact of the three interactions: abiotic–biotic interaction, within-trophic-level competitive interaction and between-trophic-level herbivory and pollination interactions during the ecological and evolutionary process. To this end, we look into how the interactive factors in the three main interactions (Figure 1) affect the population dynamics, coevolution and diversity in an ecosystem context. For the three interactions, (1) we use abiotic– biotic interaction related growth and death rates (*r*_A_, *r*_B,_ *d*_A_, *d*_B_) to represent the strength of interactions between abiotic environment and biotic populations (herbivore and pollinator populations). A population with a higher growth rate or a lower death rate means that the population has a higher positive interaction with the environment. (2) We use the competitive interaction related carrying capacities (*K*_A_, *K*_B_) to represent the strength or intensity of competitive interaction within the herbivore populations or the pollinator populations. A higher carrying capacity inside the herbivore populations or the pollinator populations means that the negative competitive interaction within focused populations is less intense. (3) Finally, we use the herbivory and pollination interaction related conversion coefficients (*c*_A_, *c*_RA_, *c*_B_, *c*_RB_) to represent the strength of the insect-plant interactions between the herbivores or the pollinators with their communal plant recourses. A higher conversion coefficient between a population and the plant resources means that positive plant-resource-based herbivory or pollination interaction is higher for that population. In the research, we examine how these factors relating to the three interactions can affect the change of ecological population sizes for both the herbivores and the pollinators. We also study in what direction the functioning traits evolve across time, that is whether the focused traits evolve to be more interactive with abiotic environment or with the biological populations in the ecosystem.

Moreover, we investigate the impact of interactive niche widths in the three interactions on population diversity. In specific, (1) we use the adaption niche widths (σ_0a_, σ_0b_) to represent the interaction range between the biological populations and abiotic environment. A higher adaptive niche width means that the trait-based population is adaptive to a wider range of environmental conditions and less sensitive to environmental change. (2) We use the competition niche widths (σ_ca_, σ_cb_) to represent the competition range within the herbivore populations or the pollinator populations. Lower competitive niche widths mean that the competition interaction ranges of the focused populations, i.e., herbivore populations (or pollinator populations), are narrow and so the herbivore population is less likely to have the overlapping living niche with other herbivore populations (or pollinator populations). In this case, the competition within the herbivore populations (or the pollinator populations) is less intense. (3) We use the resource niche widths (σ_a_, σ_b_) to represent the interactive range between the herbivore or the pollinator populations with their mutual plant resources. A higher resource niche width means that the herbivore or the pollinator populations can feed on a wide range of trait-based plant resources. Here we examine how the niche widths in the three interactions affect the diversity in the herbivore and pollinator populations.

## Results

### 1. Ecological and evolutionary responses to the three interactions

#### (1) The effect of abiotic–biotic interaction on population sizes and trait evolutions

The strength of abiotic-biotic interactions in the herbivore and pollinator populations are measured by their growth (*r*_A_, *r*_B_) and death rates (*d*_A_, *d*_B_) while living in this abiotic environment, and we explore how the antagonists and mutualists’ ecological population sizes and their evolutionary trait values can be affected the rates (*r*_A_, *r*_B_, *d*_A_, *d*_B_) (Table 1, Eq. I). As expected, our analyses show that the population size of the focal populations increases with their own growth rate and decreases with their own death rate for both the herbivores and pollinators (Table 2, Figure A1). The indirect effects propagating through the ecological network to the other populations, however, are less obvious. The increase in the growth rate (or the decrease in the death rate) of herbivores decreases the population sizes of the pollinators, due to the herbivores’ direct negative effects on the plants as the herbivores consume the plants and so also brings subsequent negative indirect effects on pollinator populations that live mutualistically with the plants. On the other hand, the increase in the growth rate (or the decrease in the death rate) of the pollinators, increases the population sizes of the herbivores, due to the pollinators’ direct positive effects on the plants and so also brings subsequent positive indirect contribution to the herbivore populations (Table 2, Figure A1 - note that the corresponding descriptive results are shown in Table 2, and the detailed results for the responses of population sizes and trait evolution to different interactive coefficient factors are shown in Appendix Figures A1).

**Table 1.** The mathematical terms in the model.

|  | Resources<br>(plants) | Antagonists<br>(herbivores) | Mutualists<br>(pollinators) |
| --- | --- | --- | --- |
| Population sizes | $R$ | $A_i$ | $B_l$ |
| Trait values | | $a_i$ | $b_l$ |
| Intrinsic growth rates | $r$ | $r_A$ | $r_B$ |
| Intrinsic death rates | $d$ | $d_A$ | $d_B$ |
| Carrying capacities | $K$ | $K_A$ | $K_B$ |
| Optimal adaptive trait values | | $a_0$ | $b_0$ |
| Optimal interactive trait values | | $a_m$ | $b_m$ |
| Adaptive niche widths | | $\sigma_{0a}$ | $\sigma_{0b}$ |
| Competitive niche width | | $\sigma_{ca}$ | $\sigma_{cb}$ |
| Resource niche width | | $\sigma_a$ | $\sigma_b$ |
| Interactive conversion coefficients | $c_{RA}, c_{RB}$ | $c_A$ | $c_B$ |

**Table 2.**
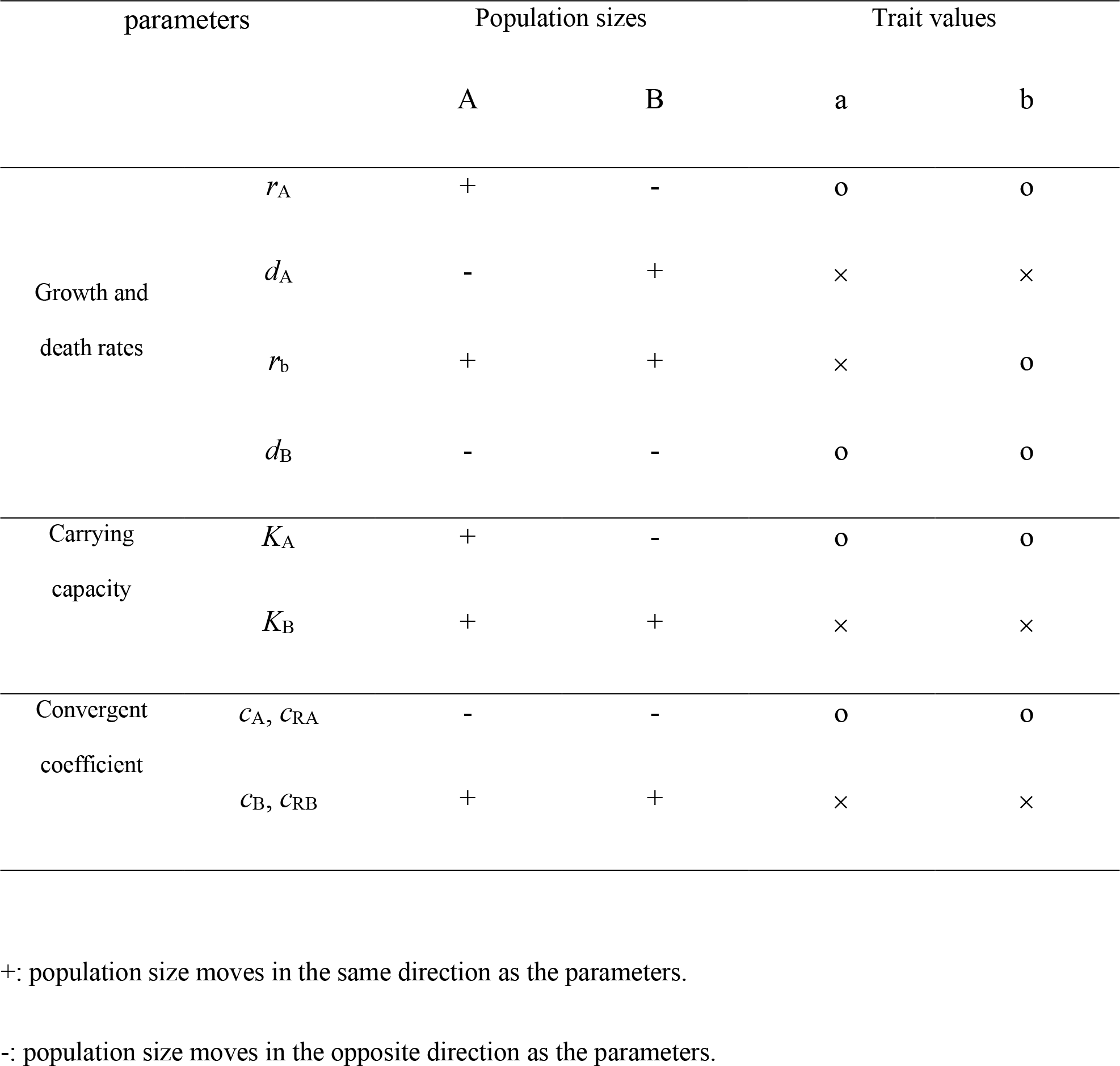
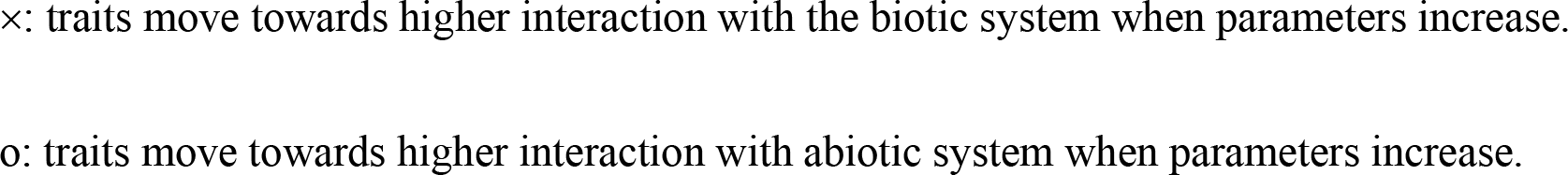
The ecological and evolutionary responses to the interactive coefficient factors.

| parameters |  | Population sizes |  | Trait values |  |
| --- | --- | --- | --- | --- | --- |
|  |  | A | B | a | b |
| Growth and death rates | $r_A$ | + | - | o | o |
| | $d_A$ | - | + | × | × |
| | $r_b$ | + | + | × | o |
| | $d_B$ | - | - | o | o |
| Carrying capacity | $K_A$ | + | - | o | o |
| | $K_B$ | + | + | × | × |
| Convergent coefficient | $c_A, c_{RA}$ | - | - | o | o |
| | $c_B, c_{RB}$ | + | + | × | × |

+: population size moves in the same direction as the parameters.
-: population size moves in the opposite direction as the parameters.
×: traits move towards higher interaction with the biotic system when parameters increase.
o: traits move towards higher interaction with abiotic system when parameters increase.

Such indirect effect also propagates across eco-evolutionary time scales (not only in the change of ecological population sizes shown above, but also in the evolution of traits). Both the trait values of herbivores and pollinators evolve towards the optimal environmental adaptive traits *a*_0_=0, *b*_0_=0, and away from the optimal interactive traits *a*_m_=1, *b*_m_=2, when the birth rate of the antagonistic herbivores (*r*_A_) increases. This means that the whole system evolves to be less interactive with other biotic populations in the system both for the pollinators and the herbivores as the antagonists adapt better to the environment. Interestingly, when the birth rate of the mutualistic pollinators (*r*_B_) increases, the pollinator traits evolve towards the optimal environmental adaptive traits and away from the optimal interactive traits. However, during such conditions, the herbivore traits evolve towards the optimal interactive traits and away from the optimal environmental adaptive trait. This result means that the pollinators evolve to rely more on abiotic environment to survive since they already have a high growth rate from the environment, whereas the herbivores evolve to rely more on the interactions with the biotic system to get more benefits while indirectly interacting with high-growth-rate pollinator populations through the ecological net (Table 2, Figure A2). As for the death rate, both the trait values of herbivores and pollinators evolve towards the optimal interactive traits *a*_m_=1, *b*_m_=2, and away from the optimal environmental adaptive traits a_0_=0, b_0_=0, when the death rate of the herbivores (*d*_A_) increases. This means that the whole system evolves to be more interactive with other biotic populations both in the pollinators and in the herbivores, when the antagonistic herbivores suffer from higher death rates. On the other hand, when the death rate of the pollinators (*d*_B_) increases, both the trait values of the herbivores and pollinators evolve away from the optimal interactive traits with the biotic system, and towards the optimal environmental adaptive traits. Thus the whole system evolves to be less interactive with other biotic populations when the mutualistic pollinators suffer from higher death rates (Table 2, Figure A2).

+: population size moves in the same direction as the parameters.

-: population size moves in the opposite direction as the parameters.

×: traits move towards higher interaction with the biotic system when parameters increase.

o: traits move towards higher interaction with abiotic system when parameters increase.

#### (2) The effect of within-trophic-level competitive interaction on population sizes and trait evolutions

The strength of within-trophic-level competitive interactions in the herbivore or pollinator populations is measured by their total carrying capacity (*K*_A_, *K*_B_) that is available for the herbivore populations or the pollinator populations to live. Higher carrying capacity means that the competitive interaction is less intense inside the herbivore populations or pollinator populations. Here we explore how the antagonists and mutualists’ ecological population sizes and their evolutionary trait values can be affected by the carrying capacity (*K*_A_, *K*_B_) (Table 1, Eq. I). As expected, the population size of either the herbivores or the pollinators increases as their own carrying capacities increase. Furthermore, since the indirect effects propagate through the network, the increase in the carrying capacity of herbivores indirectly decreases the population sizes of the pollinators. And the increase in the carrying capacity of the pollinators, on the other hand, indirectly increases the population sizes of the herbivores (Table 2, Figure A1). This result means that a lower intensity of competitive interaction within the herbivores, amplifies the herbivores’ negative effects on the whole system, so decreases the population size of the pollinators; whereas the lower intensity of competitive interaction within the pollinators, amplifies the pollinators’ positive effects on the whole system, so increases the population size of the herbivores.

Such indirect effect also propagates across eco-evolutionary time scales and shows in the evolution of traits of both herbivore and pollinator populations. Both the trait values of herbivores and pollinators evolve towards the optimal interactive traits *a*_m_=1, *b*_m_=2, and away from the optimal environmental adaptive traits *a*_0_=0, *b*_0_=0, when the carrying capacity of the pollinators (*K*_B_) increases. This means that the whole system evolves to be more interactive with other populations both in the pollinators and in the herbivores, when the mutualistic pollinators face less negative competitive interactions in the system. On the other hand, when the carrying capacity of the herbivores (*K*_A_) increases, both traits evolve away from the optimal interactive traits, and towards the optimal environmental adaptive traits. This means that the whole system evolves to rely more on abiotic environment and is less interactive with other populations, when the antagonistic herbivores face less negative competitive interactions. (Table 2, Figure A2).

#### (3) The effect of between-trophic-level herbivory and mutualistic interactions with plant resources on population sizes and trait evolutions

The strength of between-trophic-level antagonistic or mutualistic interactions is measured by the conversion coefficients when the herbivore or pollinator populations interact with the shared plant resources (*c*_A_, *c*_RA_, *c*_B_, *c*_RB_). High conversion coefficients mean that the antagonistic or mutualist interactions are stronger between the shared plant resources and the corresponding antagonists or the mutualists. Here we explore how the antagonists and mutualists’ ecological population sizes and their evolutionary trait values can be affected by the conversion coefficients (*c*_A_, *c*_RA_, *c*_B_, *c*_RB_) (Table 1, Eq. I). It is shown that when the conversion coefficients of the antagonistic herbivory interaction (*c*_A_, *c*_RA_) increase, both the herbivore (direct effect from the antagonistic herbivore interactions themselves) and the pollinator (indirect effect through the network system from the herbivores to the pollinators) population sizes decrease, because the negative antagonistic interaction in the system is higher. However, when the conversion coefficients of the mutualistic pollination interaction (*c*_B_, *c*_RB_) increase, both the herbivores (indirect effect) and the pollinators (direct effect) population sizes increase, because the positive mutualistic interaction in the system is higher (Table 2, Figure A1).

Similarly, both the direct and indirect effects propagate across eco-evolutionary time scales and also show in the evolution of traits of both herbivore and pollinator populations. When the conversion coefficients of the herbivores (*c*_A_, *c*_RA_) increase, both the trait values in the herbivores and pollinators evolve away from the optimal interactive traits with the biotic system but towards the optimal environmental adaptive traits (which means less interaction with the other biotic populations in the system, but more with abiotic environment), since the level of negative antagonistic interaction in the system is higher. However, when the conversion coefficients of the pollinators (*c*_B_, *c*_RB_) increase, both the trait values in the herbivores and pollinators evolve towards the optimal interactive traits but away from the optimal environmental adaptive traits (which means more interaction with the other biotic populations in the system, but less with abiotic environment), since the level of positive mutualistic interaction in the system is higher (Table 2, Figure A2).

### 2. The population diversity in response to the three interactions

Here we study how the trait diversity in the evolutionary time scale is affected by the interactive niche widths in the three interactions. The interactive niche widths of the herbivore and pollinator populations in the three interactions (adaptive niche width σ_0a_, σ_0b_, competitive niche width σ_0a_, σ_0b_, resource niche width σ_0a_, σ_0b_, see Figure 1 and Table 1), determine how their interactive trait mismatch can affect the growths of herbivore and pollinator populations in the dynamic ecosystem. Higher interactive niche widths mean that the corresponding interactive niches are wider and so a higher mismatch of trait values is allowed for the same growth rates. With different levels of interactive niche widths in the three interactions, both the herbivore and pollinator populations evolve and also diverge (a new population with a new distinct trait emerges) differently across evolutionary time scales. We explore how the numbers of herbivore and pollinator populations with distinct traits (as the indicator of diversity) respond to the changes of these interactive niche widths when all the populations in the ecological system reach their highest fitness in the fitness landscape in the eco-evolutionary dynamics, and so the populations no longer evolve or diverge (the *ESS*, see the Method session).

Firstly, our results show that when the adaptive niche width of herbivores (σ_0a_) or the pollinators (σ_0b_) is wider, the level of diversity is higher for the herbivore or the pollinator populations (Figure 2). This corresponds to circumstances when a broader range of abiotic environmental opportunities (e.g., living habitats, geographic location, external food resource, etc) are allowed for the herbivore or the pollinator populations to adapt to, in which case more diversified populations with distinct traits adapting to specific environmental conditions emerge in the system. Secondly, our results show that when within-trophic-level competitive niche width of herbivores (σ_ca_) or the pollinators (σ_cb_) is narrower, the level of diversity is higher for the herbivore or the pollinator populations (Figure 2). This means that the within-trophic-level competition is less intense with less overlapping conflicting living space for the herbivore or pollinator populations, then diversity is higher in the system. Thirdly, we find that when between-trophic-level resource niche width of the herbivores (μ_a_) or the pollinators (μ_b_) is wider, the level of diversity is higher for the herbivore or the pollinator populations (Figure 2). This is when the herbivore populations and the mutualist populations can benefit from a broader range of plant resources in the system, and so more diversified populations with distinct traits emerge and thrive in the system.

**Figure 2.**
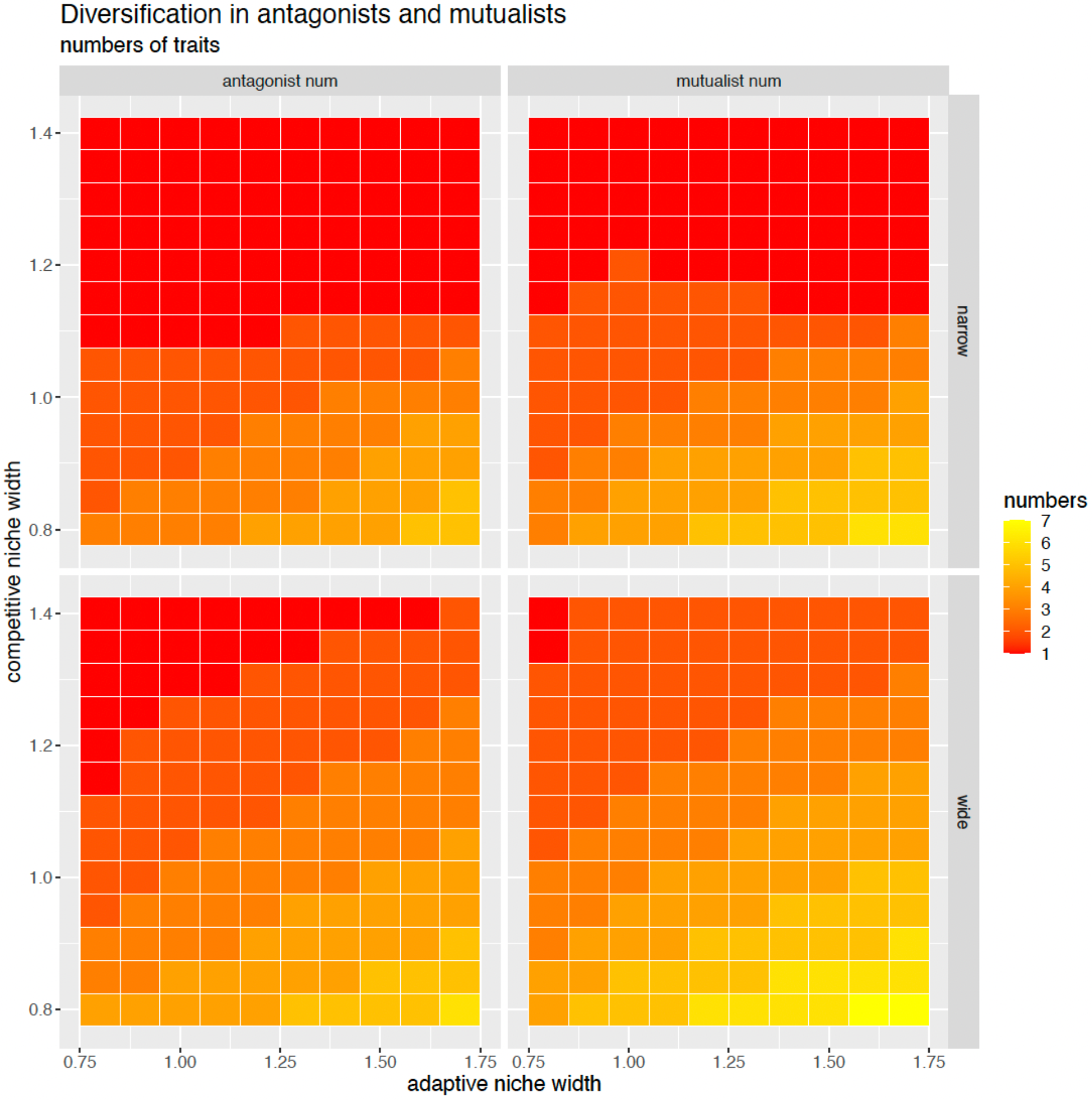
The patterns that the interactive niche width affect the diversity (the number of populations with different traits) of antagonists and mutualists. The vertical axis of each panel shows the gradient of the competitive niche widths. All four panels show that the lower the competitive niche widths the higher the diversity in traits. The horizontal axis of each panel shows the gradient of the adaptive niche width. All four panels show that the higher the adaptive niche width, the higher the diversity in traits. The upper two panels show the narrow antagonistic interactive resource niche widths (between the antagonists and plants) and the narrow mutualistic interactive resource niche widths (between the mutualists and plants). The lower two panels show the corresponding wide antagonistic and mutualistic interactive resource niche widths. Comparing the upper two with the lower two panels, the wider interactive resource niche widths lead to higher diversity for both antagonists and mutualists. The parameter values: *a*_0_=0, *b*_0_=0, *a*_m_=1, *b*_m_=2, *r*_A_=0.5, *r*_B_=0.5, *r*=1, *d*_A_=0.1, *d*_B_=0.1, *d*=0, *K*=10, *K*_A_=3, *K*_B_=3, *c*_RA_=0.2, *c*_RB_=0.2, *c*_A_=0.1, *c*_B_=0.1, σ_0a_, σ_ca_, σ_0b_, σ_cb_ equal the values shown in the figures, and σ_a_=1, σ_b_=1 (upper left, upper right), σ_a_=1.3, σ_b_=1 (lower left), σ_a_=1, σ_b_=1.3 (lower right). The parameter terms are explained in Table 1.

Notably, all these observations show a fundamental pattern: populations of herbivores or pollinators have a higher level of diversify under mild living conditions, where adaptability to the environment is high, within-trophic-level competition is lower, and herbivory or mutualistic resource availability is abundant. In contrast, harsh living conditions, characterized by narrow adaptive niches, intense within-trophic-level competition, and limited between-trophic-level resource opportunities, stifle population diversification. Note that all these differences do not affect the diversity in the other population (not shown in Figure 2).

## Discussion

In biological ecosystems, populations interact with a multitude of biotic entities, both horizontally (e.g., competitors) and vertically (e.g., trophic-level predators, herbivores, mutualists, etc), as well as interact with their abiotic surroundings, including habitats, geographic locations, and climate. To unravel this intricate web of interactions and simplify our understanding of the ecological and evolutionary processes, we identify three pivotal types of interactions that nearly all biological populations face in ecosystems: abiotic-biotic interactions^(1)^, within-trophic-level competition^(2)^, and between-trophic-level antagonistic or mutualistic interactions with the mutual plant resources^(3)^ (note that superscription (1)(2)(3) referred to the three interactions, illustrated in Figure 1, same as below). We developed a trait-based eco-evolutionary model (Eq. I) to scrutinize the responses elicited by the three interactions within an ecosystem, encompassing ecological dynamics, coevolutionary patterns, and diversity phenomena. The general trait-based nature of the model enables its use in various eco-evolutionary processes in diverse dynamic systems, including but not limited to competition, cooperation, mutualism, herbivory, predation, and abiotic-biotic interactions.

From our analyses, we find that the population sizes of both the mutualists and antagonists are shaped by their three interactions (abiotic-biotic interaction^(1)^, within-trophic-level competition^(2)^, between-trophic-level interaction with mutualists and antagonists’ shared resources^(3)^) (Figure 1) either directly from mutualists or antagonists themselves or indirectly from the other. The stronger positive mutualists’ interaction in all the three interactions, whether with abiotic environment^(1)^ (i.e., higher adaptivity to the environment) (Memmott., 2007), or among themselves^(2)^ (reduced competition), or with plant resources^(3)^ (higher compatibility with the plant resources), lead to increased population sizes for both the mutualists themselves (directly) and the antagonists (indirectly). This rise is attributed to the positive influence of mutualists on the entire ecosystem. For the antagonists, stronger interactions with abiotic environment^(1)^ or among themselves^(2)^ boost their population sizes (directly) but decrease mutualists’ population size (indirectly), due to the amplified negative impact on the shared plant resources. Antagonists’ intensified negative interaction with communal plant resources^(3)^, such as consuming plants, results in reduced population sizes for both the antagonists’ and mutualists, due to their higher detrimental impact on the communal plant resources. Other studies such as the empirical evidence, demonstrated by Fiegna & Velicer (2005), support this indirect effect in the ecological network, where cooperative populations can experience negative effects on reproduction and even face extinction when antagonistic interactions in the system are high.

The evolution of the trait values is also shaped by the strength of the three interactions by the mutualists or antagonists both directly and indirectly. The traits of antagonists evolve to be more interactive with biotic systems and less reliant on abiotic environment, when the mutualists exhibit stronger interactions with the system (either being less competitive^(2)^ or more compatible with plant resources^(3)^ or the environment^(1)^), or when the antagonists exhibit weaker corresponding interactions with the system. This signifies the significantly positive contributions of mutualists to the system, contrasting with the negative contributions brought by antagonists. On the other hand, for the mutualists, strong positive interaction with the biotic populations (reduced competition^(2)^ or higher compatibility with the plant resources^(3)^), drives their traits to become more interactive with the biotic system than with abiotic system. However, when their interaction with abiotic environment^(1)^ is stronger, their traits evolve to be more interactive with abiotic system. The mutualists’ traits also evolve to be less interactive with the biotic system and more interactive with abiotic system when any of the three interactions with antagonists are stronger. To sum up, the mutualists evolve to interact more with the biotic system when they gain more from their mutualist relation with the biotic system, but evolve to interact less when they are more exploited by the negative antagonistic or competitive relation in the biotic system. Meanwhile, antagonists evolve to interact more with the biotic system when they benefit from mutualists’ positive contribution to the system, yet evolve to interact less with the biotic system when their own negative antagonistic relation within the system intensifies.

Diversity among both antagonists and mutualists is also significantly influenced by all the three interactions and mainly through their interactive niche width in each interaction (Leibold, M. A., 1995). Our finding shows that a narrower competitive niche width^(2)^, indicating a reduced intensity of competition among populations, correlates with higher diversity. This finding aligns with the theory suggesting that reduced competition intensity within populations, resulting from a narrower overlap in functional traits, creates more opportunities for distinct adaptations, and therefore higher diversified interactive populations emerge (Dieckmann & Doebeli, 1999; Doebeli & Dieckmann, 2000; Pontarp et al., 2015; Pontarp et al., 2017; Pontarp et al., 2012).

Our research also reveals that a wider adaptive niche width^(1)^, indicating populations’ greater adaptability to a broader range of environments (reflecting higher ecological opportunity), correlates with increased diversity. This aligns with other studies (Petanidou & Potts., 2006; Wellborn & Langerhans, 2015; Wiens & Graham., 2005) that a wide range of ecological opportunities provides varied distinct niches in habitat and other living resources, so that different species can evolve to exploit their own specific niches, and thus adaptive radiation and coexistence of species can happen. This phenomenon is often observed in isolated environments, such as islands or newly formed habitats, where diverse ecological niches are available for colonization (Grant & Grant., 2008; Losos., 2011; Thorpe & Malhotra., 1996; Yoder et al., 2010). The empirical review study by Bolnick and Fitzpatrick (2007) explains that sympatric diversification often involves adaptation to different ecological niches, which can result in expended adaptive niche widths and, consequently, lead to higher diversification rates within a single environment. This suggests a further positive loop between the extension of adaptive niche width and the higher diversified ecological populations, which is worth further investigation.

Finally, our study shows that a wider trophic-level biotic resource niche width^(3)^ (e.g., here is the communal plant resource) is associated with higher diversity among herbivores or pollinators. Like the above adaptive niche width^(1)^, provided with varied living-relied plant resources, herbivores or pollinators with different specific function-based traits (associated with plant consumption or pollination) can coexist (Valdovinos et al., 2013; Vazquez & Aizen., 2006). This result is well recorded in many studies that the biodiversity in the tropics is much more abundant than the poles (Dowle et al., 2013; Hillebrand., 2004; Mora & Robertson., 2005). Also herbivore insects are found to evolve specialized feeding behaviours or physiological adaptations to exploit specific plant defences and chemical compounds (Farrell., 1998; Wang et al., 2019); and pollinators, such as hummingbirds, are shown to evolve various bill shapes and lengths to match the specific floral traits of the plants they pollinate, showing the adaptation of pollinators to different plant opportunities (Curti & Ortega-Baes., 2011; Valdovinos et al., 2013).

To sum up, our study elucidates that biological populations tend to experience higher diversity under relatively milder conditions (wider abiotic environmental opportunity^(1)^, reduced competition intensity^(2)^, or broader benefits from other biotic resources^(3)^), but lower diversity under relatively harsher conditions. These findings underscore the important role of niche widths in shaping the biodiversity of biological populations. Milder conditions, with their wider adaptive or interacting possibilities, both abiotic and biotic, provide biological populations with ample spaces to thrive and diversify.

The study here identifies three very main interactions that are found to be important in most of the ecological systems: between-trophic-level^(3)^ and within-trophic-level biotic^(2)^ and also abiotic-biotic interactions^(1)^. These interactions are studied in many studies mentioned above, but most of them are confined to one specific interaction. Here we have a comprehensive study to address all these three interactions in concert with both antagonistic and mutualist contexts and try to build a theoretical foundation to study the overall ecological network. We covered some common themes such as mutualism, antagonism, competition, and adaptation to elucidate the underlying reasons for population structure changes, trait evolution, and biodiversity in the interactive dynamic ecosystem. We hope broad implications can be derived from these common themes and the related mechanisms from this study. For example, several pressing and intriguing questions are often discussed - why certain species face extinction or experience significant population declines due to their currently changing abiotic or biotic interactions; why certain species alter their habitats or traits, or coevolve dynamically with other species; why the biodiversity in certain areas is much higher than others (e.g., areas with different latitude or elevation); how can we preserve endangered species amidst challenging environmental or climate conditions; how can we explain the loss of biodiversity and explore avenues to prevent further deterioration of these situations. We hope that our theoretical analyses provide the preliminary foundation to address these critical issues, and also aid in future predictions and interventions.

## Appendix

**Figure A1.**
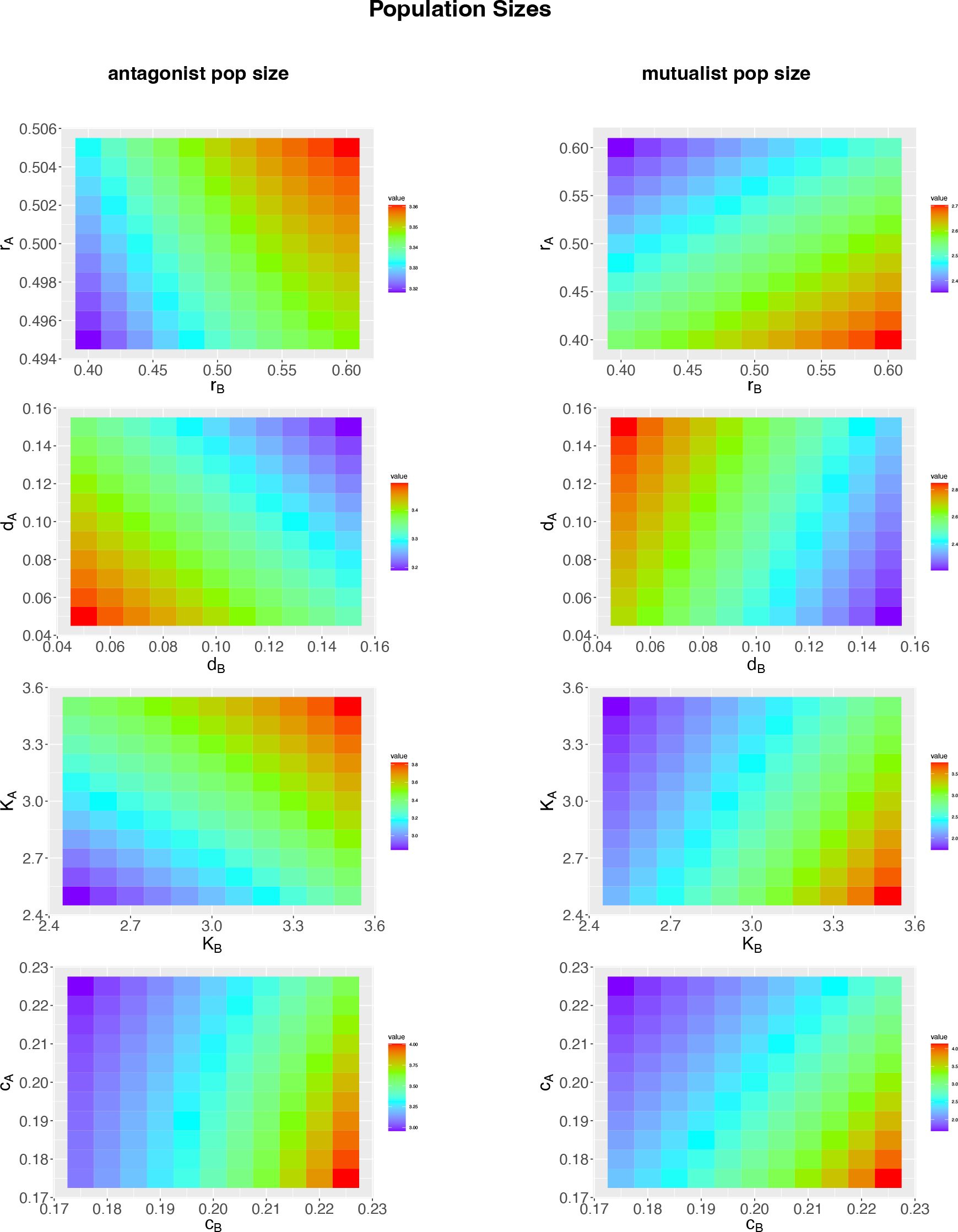
The responses of population sizes of antagonists (left) and mutualists (right) to the interaction factors (grow rates, death rates, carrying capacity, convergent coefficients in the 1^st^, (2)^nd^, 3^rd^, 4^th^ rows correspondingly). The parameter values: *a*_0_=0, *b*_0_=0, *a*_m_=1, *b*_m_=2, *r*=1, *d*=0, *K*=10, σ_0a_= σ_ca_= σ_a_=1, σ_0b_= σ_cb_= σ_b_=1, *r*_A_, *r*_B_, *d*_A_, *d*_B_, *K*_A_, *K*_B_, *c*_RA_, *c*_RB_, *c*_A_=0.5*c*_RA_, *c*_B_=0.5*c*_RB_ (here shows for *c*_A_ and *c*_B_ and omits *c*_RA_ and *c*_RB_) equal the values shown in the figures. The parameter terms are explained in Table 1.

**Figure A2.**
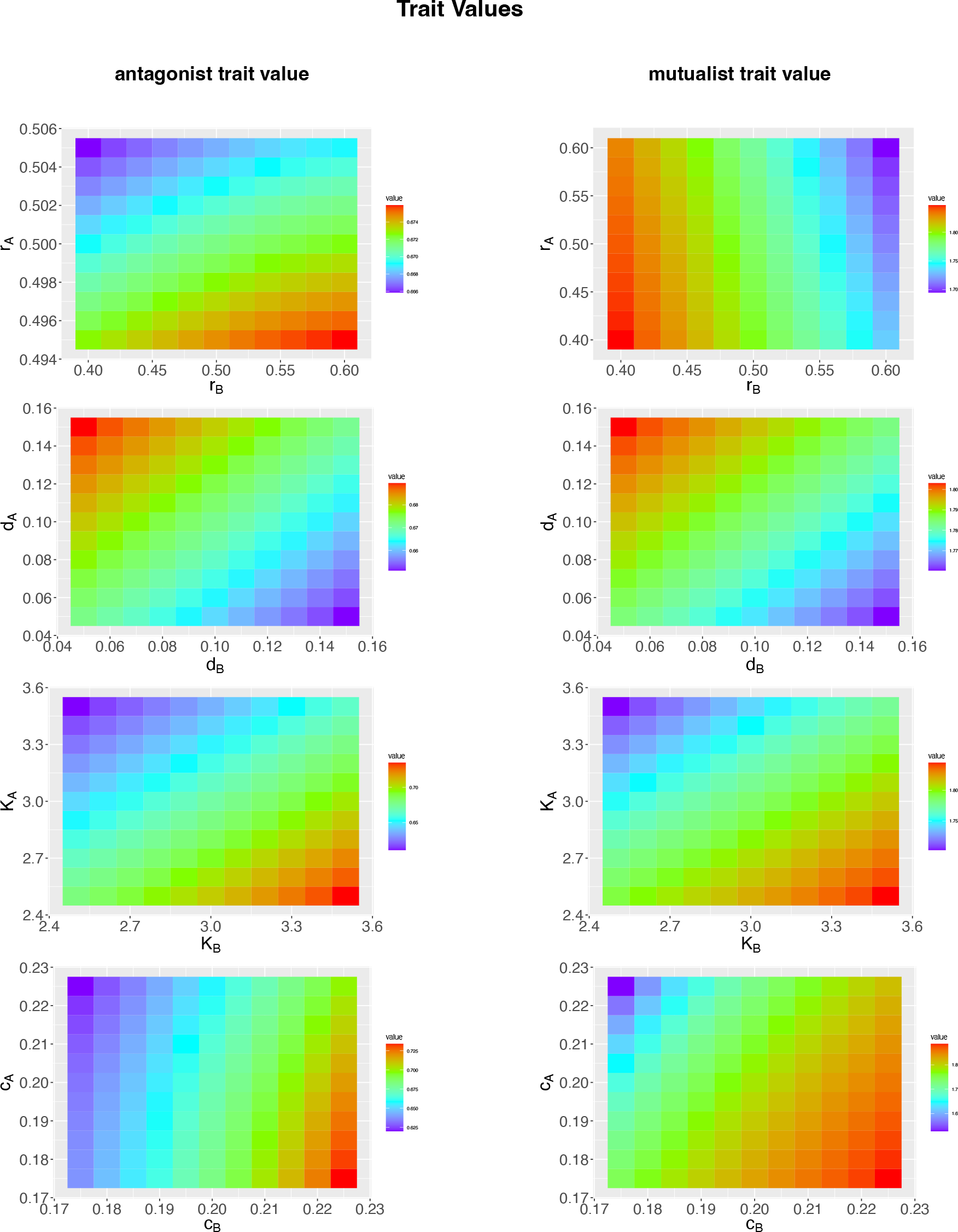
The responses of trait evolution of antagonists (left) and mutualists (right) to the interaction factors (growth rates, death rates, carrying capacity, and convergent coefficients in the 1^st^, 2^nd^, 3^rd^, 4^th^ rows correspondingly). The parameter values: *a*_0_=0, *b*_0_=0, *a*_m_=1, *b*_m_=2, *r*=1, *d*=0, *K*=10, σ_0a_= σ_ca_= σ_a_=1, σ_0b_= σ_cb_= σ_b_=1, *r*_A_, *r*_B_, *d*_A_, *d*_B_, *K*_A_, *K*_B_, *c*_RA_, *c*_RB_, *c*_A_=0.5*c*_RA_, *c*_B_=0.5*c*_RB_ equal the values shown in the figures. The parameter terms are explained in Table 1.

